# Transient Reduction of DNA Methylation at the Onset of Meiosis in Male Mice

**DOI:** 10.1101/177535

**Authors:** Valeriya Gaysinskaya, Brendan F. Miller, Godfried W. van der Heijden, Kasper D. Hansen, Alex Bortvin

**Affiliations:** Department of Embryology, Carnegie Institution for Science, Baltimore, Maryland, United States of America; Department of Biology, Johns Hopkins University, Baltimore, Maryland, United States of America; Translational and Functional Genomics Branch, National Human Genome Research Institute, National Institutes of Health, Bethesda, Maryland, United States of America; Department of Obstetrics and Gynaecology, Erasmus MC, University Medical Center, PO BOX 2040, 3000 CA, Rotterdam, The Netherlands; Department of Biostatistics, Johns Hopkins University, Baltimore, Maryland, United States of America; Center for Computational Biology, Johns Hopkins University, Baltimore, Maryland, United States of America; McKusick-Nathans Institute of Genetic Medicine, Johns Hopkins University, Baltimore, Maryland, United States of America

**Keywords:** DNA methylation, meiosis, DNA replication, spermatogenesis, mouse, LINE-1

## Abstract

The quality of germ cells depends on successful chromatin organization in meiotic prophase I (MPI). To better understand the epigenetic context of MPI we studied the dynamics of DNA methylation in wild-type male mice. We discovered an extended period of genome-wide transient reduction of DNA methylation (TRDM) during early MPI. Our data show that TRDM arises by passive demethylation in the premeiotic S phase highlighting the abundance of hemimethylated DNA in MPI. Importantly, TRDM unmasks a deficit in retrotransposon LINE-1 DNA methylation contributing to its expression in early MPI. We propose that TRDM facilitates meiosis and gamete quality control.

## Introduction

Meiosis is a specialized cell cycle that yields haploid gametes. It involves a single round of premeiotic DNA replication followed by two successive meiotic divisions. Aberrant chromosome segregation in meiotic divisions causes germ cell and embryo aneuploidy (Webster and Schuh 2017). Precise chromosome segregation in meiosis and gamete quality critically depend on a protracted meiotic prophase I (MPI) when homologous chromosomes pair, synapse and recombine (Zickler and Kleckner 2015). The corresponding changes in the appearance of meiotic chromosomes allow further subdivision of MPI into preleptotene (PL), leptotene (L), zygotene (Z), pachytene (P) and diplotene (D) substages to facilitate its analysis (Supplemental Fig. S1).

Since all MPI processes occur in the chromatin context, it is not surprising that perturbations of histone modifications and DNA methyltransferases impact MPI (Crichton et al. 2014; Szekvolgyi et al. 2015; Yelina et al. 2015a; Zamudio et al. 2015). However, despite the wealth of evidence of meiotic failure of mouse male germ cells deficient in DNA methylation (Bourc’his and Bestor 2004; Webster et al. 2005; Crichton et al. 2014; Zamudio et al. 2015; Barau et al. 2016) and detailed knowledge of genome-wide DNA methylation in postnatal spermatogonia (Hammoud et al. 2015; Kubo et al. 2015), the precise dynamics of this crucial epigenetic modification during MPI remain unknown. Although bulk DNA methylation of male germ cells precedes meiosis (Oakes et al. 2007), a long-standing observation of transient expression of LINE-1 (L1) retrotransposons at the onset of meiosis, indicates DNA methylation changes in MPI (Branciforte and Martin 1994; Soper et al. 2008; van der Heijden and Bortvin 2009). Thus, in this study, we investigated the dynamics of DNA methylation across MPI and reveal a protracted period of genome-wide transient reduction of DNA methylation (TRDM) during MPI, a previously unrecognized epigenetic feature of meiotic chromosomes in male mice.

## Results and Discussion

To characterize the dynamics of DNA methylation across MPI, we used an optimized flow cytometry cell sorting method to obtain two biological replicates of spermatogonial (Spg), PL, L, Z, P, D spermatocytes and epididymal spermatozoa (Spz) (Gaysinskaya et al. 2014) (Supplemental Fig. S2). The purity of MPI cell fractions was verified by staining for meiosis-specific (SYCP3, γH2AX) and spermatogonia-enriched (DMRT1, DMRT6) markers as described previously (Gaysinskaya and Bortvin 2015). Critically, all germ cell fractions were devoid of somatic cells (Materials and Methods). Additionally, gene expression profiling of a wide panel of soma-enriched and germ cell-specific genes by RNA-seq confirmed the purity and stage-specificity of our samples (Supplemental Fig. S3). Using these samples, we performed whole-genome bisulfite DNA sequencing (WGBS) for genome-wide analysis of DNA methylation at single-CpG resolution (Supplemental Table S1). Over 90% of reads aligned to the mouse genome and exhibited high efficiency of bisulfite conversion (Supplemental Tables S1 and S2). Each biological replicate accounted for 87-94% of genomic CpGs with 3 to 6 x average of CpG coverage per individual sample after read deduplication and processing (Supplemental Table S3). Pairs of biological replicates exhibited high inter-individual Pearson correlation indicating excellent reproducibility of our data (Supplemental Table S4). Since cytosine methylation levels at non-CpG sites were negligible (0.3-0.4%), we excluded them from later analyses (Supplemental Table S4).

First, we examined genome-wide DNA methylation levels across MPI. Consistent with the notion of high DNA methylation levels in spermatogenesis, we observed high median (> 84%) levels of CpG methylation of Spg, P, D and Spz genomes (Fig. 1A, Supplemental Table S5). Interestingly, we uncovered an extended window of reduced global DNA methylation during early MPI demarcated by a pronounced drop in DNA methylation (13 pp) levels from PL and to P followed by a progressive gain of DNA methylation in L and Z, returning to premeiotic levels by P (Fig. 1A). Overall, a period of reduced global DNA methylation lasts from PL to P (∼70 hr), and we will refer to it as transient reduction of DNA methylation at the onset of meiosis (TRDM) (Fig. 1A, Supplemental Table S5).

**Fig. 1.**
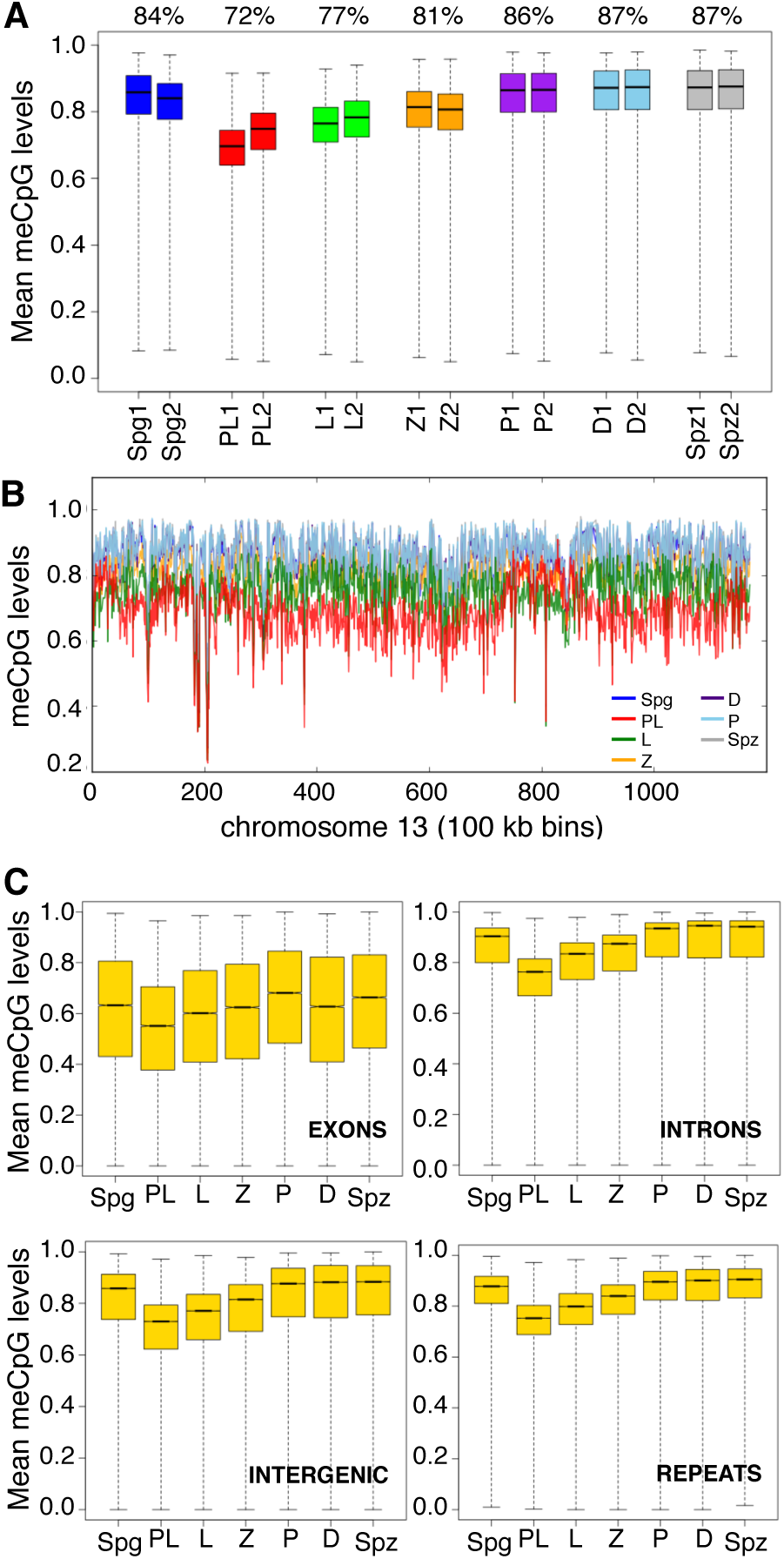
Global DNA methylation dynamics in MPI. (A) Genome-wide DNA methylation was summarized as means of non-overlapping bins of 500 CpGs for individual biological replicates. Box-and-Whisker plot shows the maximum, upper quartile, median, lower quartile and minimum of data. Median percent DNA methylation for both replicates is specified above the boxplot. (B) Chromosome-wide DNA methylation levels were plotted across chromosome length (chromosome 13, replicate 1 is shown). DNA methylation was averaged using sliding non-overlapping bins of 100 kbp. (C) Box-and-Whisker plot of DNA methylation levels across various genomic features. The average DNA methylation levels were aggregated as consecutive, non-overlapping averages of 100 CpGs. Averages were combined for biological replicates.

To examine the chromosome-wide distribution of DNA methylation in individual MPI substages, we summarized DNA methylation levels over a distance of 100 kb-wide non-overlapping windows spanning the length of each chromosome. We found that global hypomethylation in PL is chromosome-wide (Fig. 1B). This was true for all autosomes examined in both biological replicates (Supplemental Figs. S4 and S5). Interestingly, although the X chromosome also exhibits TRDM, it tends to be less methylated in all MPI substages (Supplemental Figs. S4 – S6). X chromosome DNA methylation levels in Spg-to-PL and PL-to-L transitions are distinctly less correlated than to the autosomes, further suggesting differences in the dynamics of its demethylation and remethylation (Supplemental Fig. S6). Nonetheless, these results showed that TRDM holds true for all chromosomes and that remethylation in MPI appears as a gradual chromosome-wide process.

To determine if DNA hypomethylation in PL is specific to a particular genomic feature, we examined DNA methylation dynamics of exons, introns, intergenic and repetitive regions, as well as functionally specialized sequences such as promoters and CpG islands (CGIs) (Fig. 1C, Supplemental Fig. S7A, Supplemental Table S6). This analysis showed that all genomic features were highly methylated in Spg and then demethylated in PL (most prominently at introns, repeats, and intergenic regions), except for CGIs whose methylation levels are already very low. Likewise, the analysis revealed comparable DNA methylation dynamics of repetitive DNA with major classes of TEs, namely, the LINEs, SINEs, LTRs and DNA transposons (Supplemental Fig. S7B). Finally, we asked if differentially methylated regions (DMRs) of imprinted genes also become hypomethylated in PL. The analysis of a subset of imprinted DMRs (Tomizawa et al. 2011) showed that DNA methylation levels of paternal imprinted DMRs follow the same dynamic observed for other genomic features while maternal DMRs remained unmethylated as expected (Supplemental Fig. 7C). Cumulatively, these results show that TRDM is indeed a genome-wide event that encompasses all chromosomes and all genomic features.

To better understand the DNA methylation dynamics in MPI, we identified regions that exhibited significant differences in methylation levels between any two consecutive MPI substages in a statistically-principled, coverage-conscious and biological replicate-aware manner (Hansen et al. 2012). This analysis revealed thousands of DMRs supporting the results of our genome- and chromosome-wide analyses (Fig. 2, Supplemental Tables S7 and S8A). Formation of large hypomethylated DMRs (with a median size of ∼35 kb, the median number of CpGs around 257 and implicating over half of the mouse genome) marked the Spg-to-PL transition (Fig. 2A,B). As a result of gradual remethylation of hypomethylated Spg-to-PL DMRs in L and Z, their mean methylation difference and sizes also progressively decreased (Fig.2B,C). Thus, while Spg-to-PL DMRs included ∼56% of all evaluated CpGs, PL-to-L and L-to-Z DMRs included ∼ 41% and ∼3% of all CpGs, respectively (Supplemental Table 8A). By intersecting genomic coordinates of DMRs between MPI substages we found that Spg-to-PL DMRs accounted for up to 75% of all PL-to-L and 63% of L-to-Z DMRs (Supplemental Materials and Methods). Therefore, sharp demethylation in PL is followed by gradual remethylation of the same CpGs in L and Z germ cells.

**Fig. 2.**
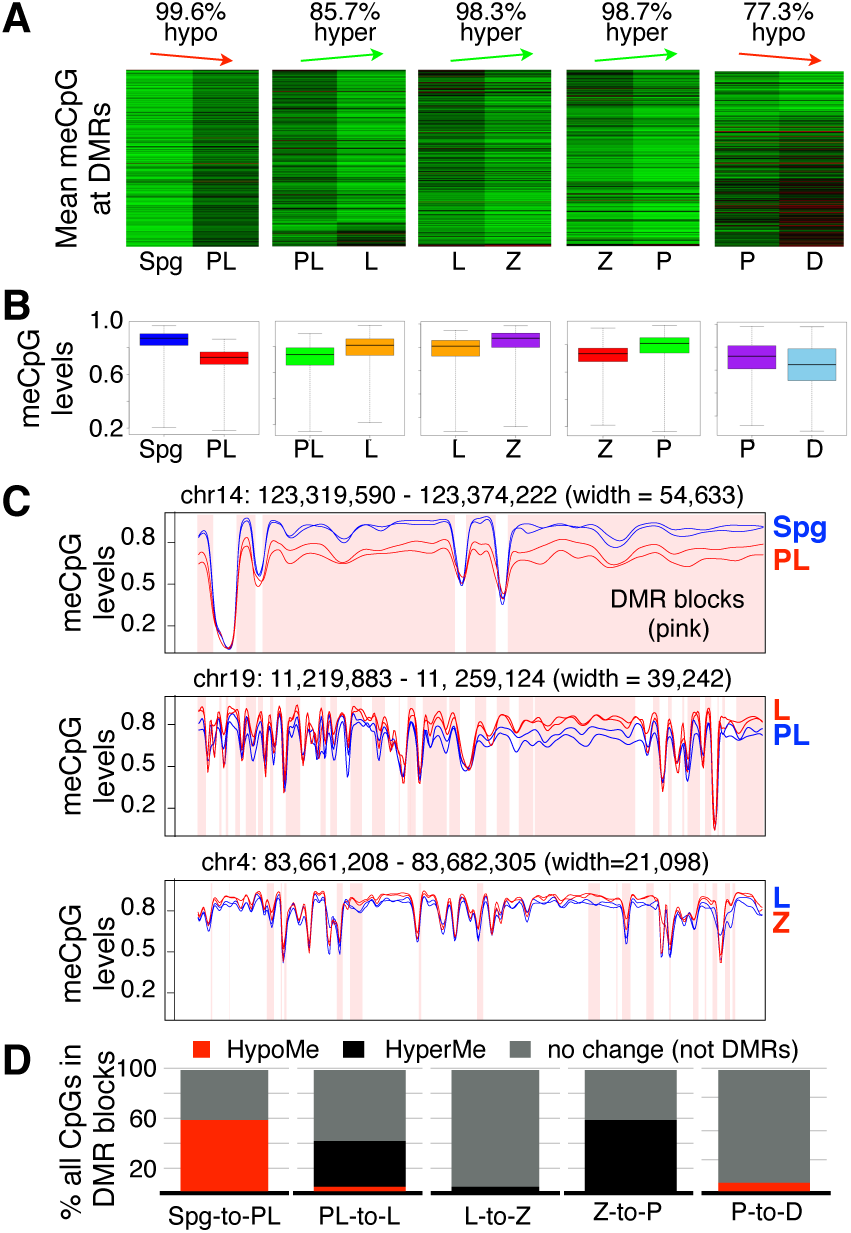
DMR dynamics in MPI. (A) Heatmap of DNA methylation profiles of all DMRs between consecutive MPI substages. Each row shows DNA methylation level of a DMR in a pairwise comparison. DNA methylation level is scaled according to red (low)-to-green (high) color scale. The percentage of DMRs exhibiting the main direction of DNA methylation change is indicated above the plots. (B) Boxplot showing DNA methylation value distribution at DMRs in MPI between two consecutive stages. (C) Smoothed DNA methylation at DMRs for Spg and PL (top), PL and L (middle), and P and D (bottom). (D) The proportion of CpGs accounted for by hypomethylated and hypermethylated DMRs.

Although global levels of DNA methylation in Z were higher relative to the preceding substages, Z is still hypomethylated relative to P (Fig. 2A). Accordingly, our DMR analysis showed that during the Z-to-P transition there is an increase in methylation at ∼57% of analyzed CpGs from 81% in Z to 88% P (Supplemental Table S8A, Fig. 2B). Therefore, while the bulk of remethylation occurs by Z, remethylation that reaches pre-meiotic or almost Spz-like levels occurs between Z and P. Indeed, the original Spg-to-PL DMRs explain most (∼75%) of all DMRs observed between Z and P (Supplemental Materials and Methods). We find that gradual remethylation concerns all genomic features examined (exons, introns coding sequences, and repeats) (Supplemental Table S8B). In Z, up to 60% of these features are still hypomethylated compared to P, although mean DNA methylation difference is relatively small (Fig. 2D, Supplemental Table S8A). In P, less than two percent of these features are found in hypomethylated DMRs relative to D, due to remethylation.

Interestingly, while P and D share very similar DNA methylation profiles overall (Fig. 1A), we observed the emergence of hypomethylated P-to-D DMRs that involve 8% of all examined CpGs in common and are of a relatively small mean genomic size (∼9 kb, as compared to ∼35 kb in PL) (Fig 2A,B,D, Supplemental Table S8A). Considering that Spg-to-PL DMRs account for only 50% of all P-to-D DMRs (Supplemental Table S8A), it is likely that the hypomethylation observed in late MPI is unrelated to TRDM.

The discovery of TRDM raised the question of its mechanistic origin. By visually inspecting patterns of chromosome-wide DNA methylation levels, we observed a possible clue to the cause of hypomethylation in PL. Focusing on PL DNA methylation trace along a chromosome, one can observe regions of relative DNA hypomethylation interrupted by a few prominent regions of relative hypermethylation (Fig. 3A, Supplemental Figs. S4 and S5). In fact, every chromosome in both PL biological replicates possessed such prominent subchromosomal domains (Supplemental Figs. S4, S5, S8 and S9A). Furthermore, the subchromosomal domains of higher relative DNA methylation levels in PL, show lower DNA methylation levels in L, resulting in an apparent switch in DNA methylation traces in these MPI substages when compared to the rest of the chromosome (Fig. 3A top panel, Supplemental Figs. S8A and S9A).

**Fig. 3.**
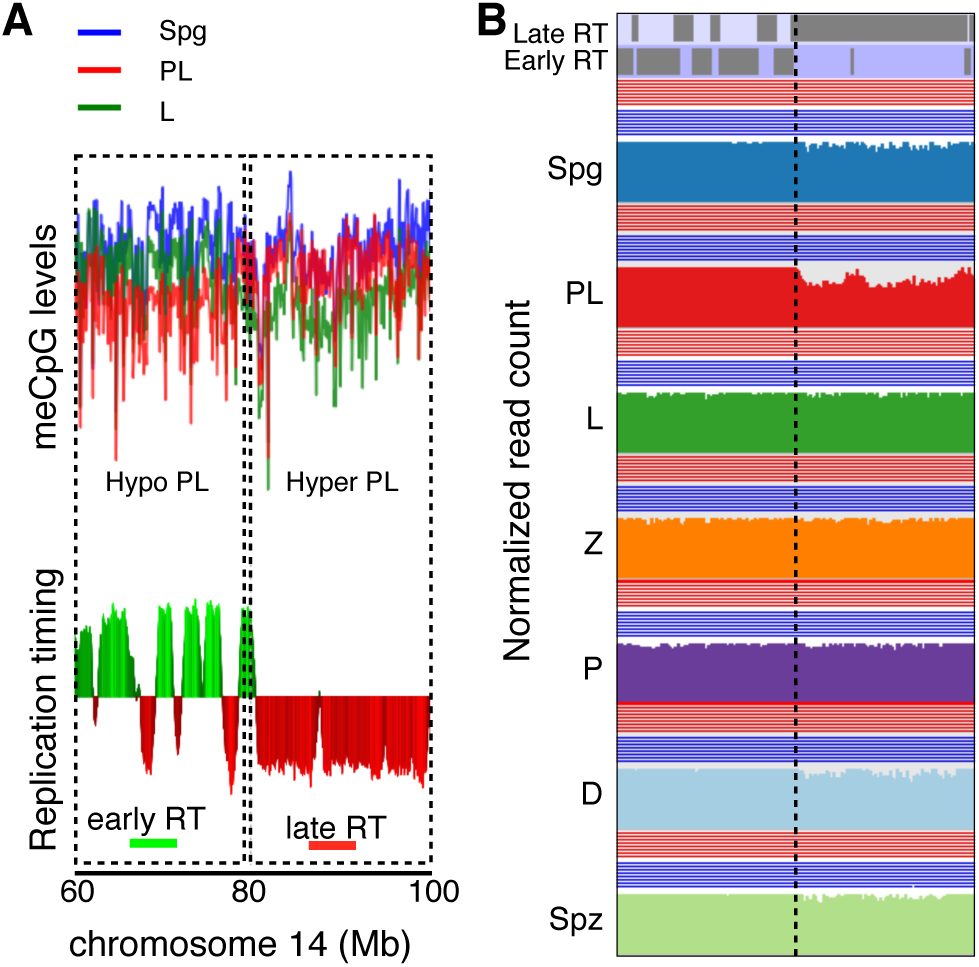
DNA methylation pattern in PL overlaps with replication timing. (A, top) The plot of CpG DNA methylation averaged using sliding non-overlapping 100 kbp windows in Spg, PL and L across a region of chromosome 14 and (A, bottom) replication timing (RT) data for the same region of chromosome 14 from mouse B-cell lymphoma CH12 cells. (B) Normalized genome sequencing coverage after WGBS summarized as averages of sliding non-overlapping 5 kbp windows.

Given the above dynamics of DNA methylation in the PL-to-L transition, we considered a role for DNA replication and replication timing domains in this phenomenon. Replication domains are large-scale genomic territories that replicate at particular times during S phase (Ryba et al. 2010; Pope et al. 2012). Global early or late replication timing profiles appear relatively preserved between different cell lines and cell types tested, although there are tissue-specific differences (Ryba et al. 2010; Yaffe et al. 2010). Remarkably, an overlay of the chromosome-wide DNA methylation pattern from our data with replication timing domains of a mouse B-cell lymphoma CH12 cell line (Weddington et al. 2008) revealed a strong overlap between the two (Figs. 3A). Specifically, in PL, we observe an overlap between large hypermethylated regions and late replicating domains. Correspondingly, an overlap is observed in PL between large-scale hypomethylated regions and early replicating domains. Interestingly, a switch between DNA hypo- and hypermethylation in PL is marked by an opposite switch in DNA methylation pattern in L (Fig. 3A). This switch in DNA methylation pattern in PL-to-L transition matched the transition from early to late replication timing domains (Fig. 3A). The overlap between DNA methylation pattern and replication timing pattern in PL was true of both biological replicates (e.g. Fig 3A, Supplemental Figs. S8, and S9). To test the strength of the association of a switch in DNA methylation levels with late replication genome-wide, we determined their Pearson correlation coefficient in the course of MPI. This analysis showed an abrupt switch in the directionality of correlation from PL to L supporting that late replicating domains switch from high to low DNA methylation levels between the two MPI stages (Supplemental Fig. S10A).

To further explore a role of DNA replication in hypomethylation of the PL genome, we evaluated the uniformity of genome sequencing coverage in our WGBS data (Fig. 3B). Previously, DNA sequence coverage was used to estimate replication timing and to evaluate underreplication in *Drosophila* polytene chromosomes (Koren et al. 2014; Yarosh and Spradling 2014). We summarized read frequency over a distance of 5kb non-overlapping windows spanning the length of the chromosome and corrected for the difference in total read count between the samples. Remarkably, we observe consistently lower sequencing coverage in the hypermethylated regions/ early replication timing domains in PL, disappearing in L (Fig. 3B, Supplemental Figs. S8 and S9). The lower sequencing coverage in PL is consistent with DNA replication during this time, while recovery of sequence coverage in L agrees with the lack of replication in L, as no replication occurs then and during the rest of meiosis. To confirm that PL spermatocytes used in our studies are replicative, we performed FACS enrichment of PL cells from mice injected with EdU 2 hrs prior to cell sorting. Subsequent EdU detection showed that >70% of FACS-enriched PL cells were replicative, with the majority of EdU patterns corresponding to middle and late S phase (Supplemental Fig. S10B) (Boateng et al. 2013).

The above results suggested that DNA replication in PL dilutes DNA methylation levels by creating hemimethylated DNA. To test this possibility directly, we analyzed methylation of complementary DNA strands using hairpin-bisulfite analysis (Arand et al. 2012) combined with next generation sequencing (Supplemental Materials and Methods). Here we focused on the 5‵-end sequence of full-length L1 elements in L1MdTf_I and L1Md_Tf_II families (Sookdeo et al. 2013). The mouse genome has ∼3000 of such elements thus permitting simultaneous measurement of DNA hemimethylation throughout the genome. After PCR amplification of hairpin-bisulfite products using L1MdTf-specific primers (Supplemental Fig. S11) followed by Illumina paired end sequencing of these L1MdTf amplicons, we obtained a range of x-y reads per each analyzed MPI stage (Supplemental Table S9). We reliably determined methylation states of 5 L1 CpG dyads (Supplemental Fig. S11) before, during and after MPI. This analysis revealed a robust increase in L1 CpG hemimethylation in PL that was followed by gradual remethylation in MPI (Fig. 4A,B). Notably, L1 hemimethylation levels and dynamics in MPI paralleled those of LINE elements in our genome-wide DNA methylation studies (compare Fig. 4A, red values, with Supplemental Fig. S7B). Importantly, by excluding hemimethylated L1 DNA from this analysis we effectively erased TRDM of L1 (Fig. 4B, blue values). Together with above evidence, these results strongly support the primary mechanism of TRDM by DNA replication-coupled passive DNA demethylation. Finally, this analysis also revealed the emergence of L1s bearing no methylation on analyzed CpG (Fig. 4C) suggesting that TRDM unmasks incompletely methylated L1 elements thus providing an opportunity for their expression at the onset of MPI.

**Fig. 4.**
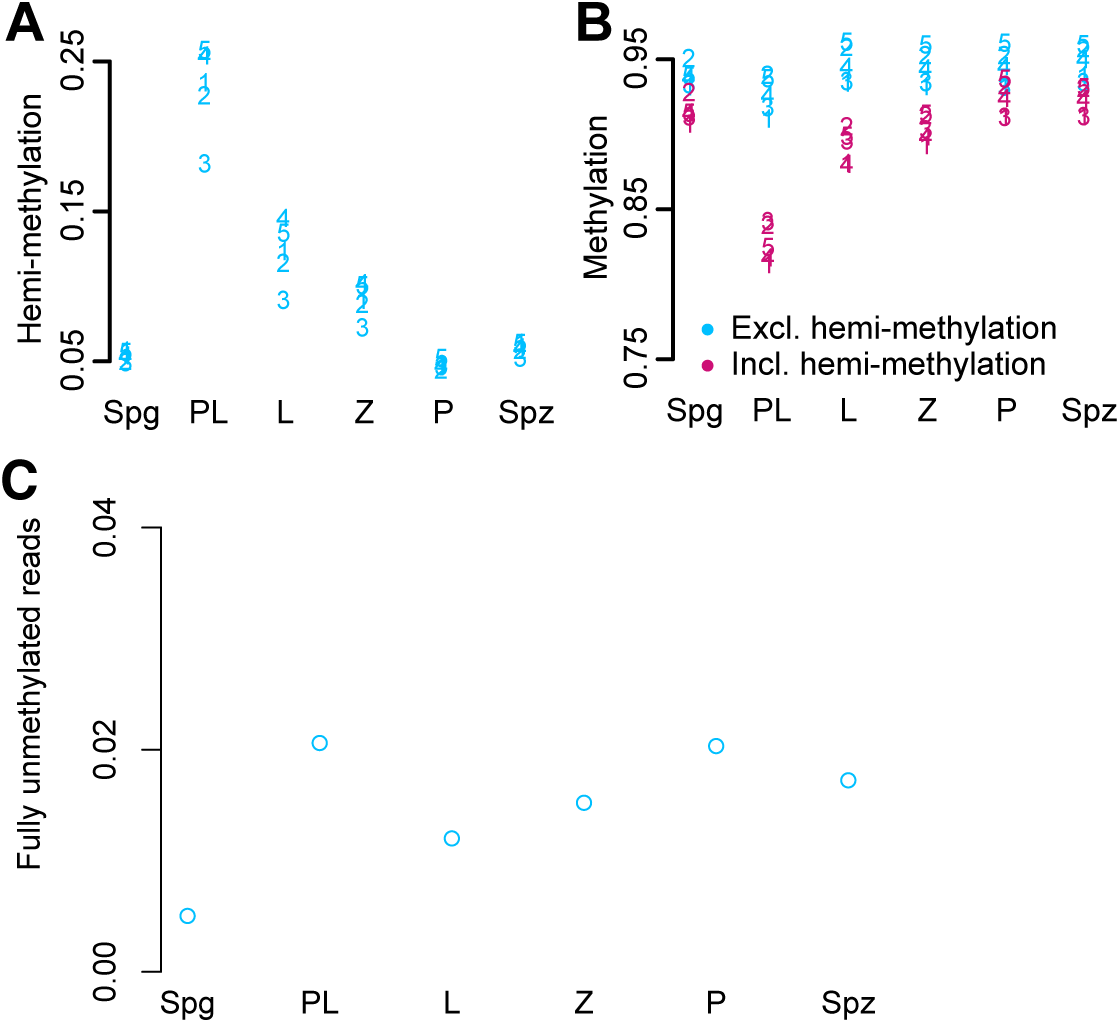
Hemimethylation of L1. Hemimethylated, methylated and unmethylated levels were quantified at 5 CpGs in L1. (A) The proportion of reads supporting hemimethylation at each of the 5 CpGs in 6 different cell states. (B) The amount of methylation at the 5 CpGs quantified including hemimethylation (pink) and excluding hemimethylation (blue). (C). Levels of fully unmethylated L1 elements in MPI based on reads where all 5 CpGs are unmethylated on both strands.

To test if TRDM of potentially active L1s in PL contributes to their expression in MPI in wild-type animals, we performed RNA-seq of FACS enriched individual MPI cell populations (Supplemental Materials and Methods). To analyze RNA abundance of TEs, we used RepEnrich strategy to account for most TE-derived RNA by way of counting both uniquely mapped and multi-mapped reads in our RNA-seq data (Criscione et al. 2014). Using this strategy, we found that transcript abundance for repeat elements as a whole shows an overall decrease from Spg onwards, with lowest levels in Spz (Supplemental Fig. S12). Intriguingly, we find that Spg-to-PL and PL-to-L transitions are accompanied by transcriptional upregulation of many classes of LINE elements (Fig. 5A). This upregulation includes all classes of potentially active L1 elements, whose expression begins to decrease in Z and is essentially extinguished by P (Fig. 5B). Interestingly, a P-to-D transition involves a strong upregulation of LINEs including potentially active LINE members (Fig. 5B).

**Fig. 5.**
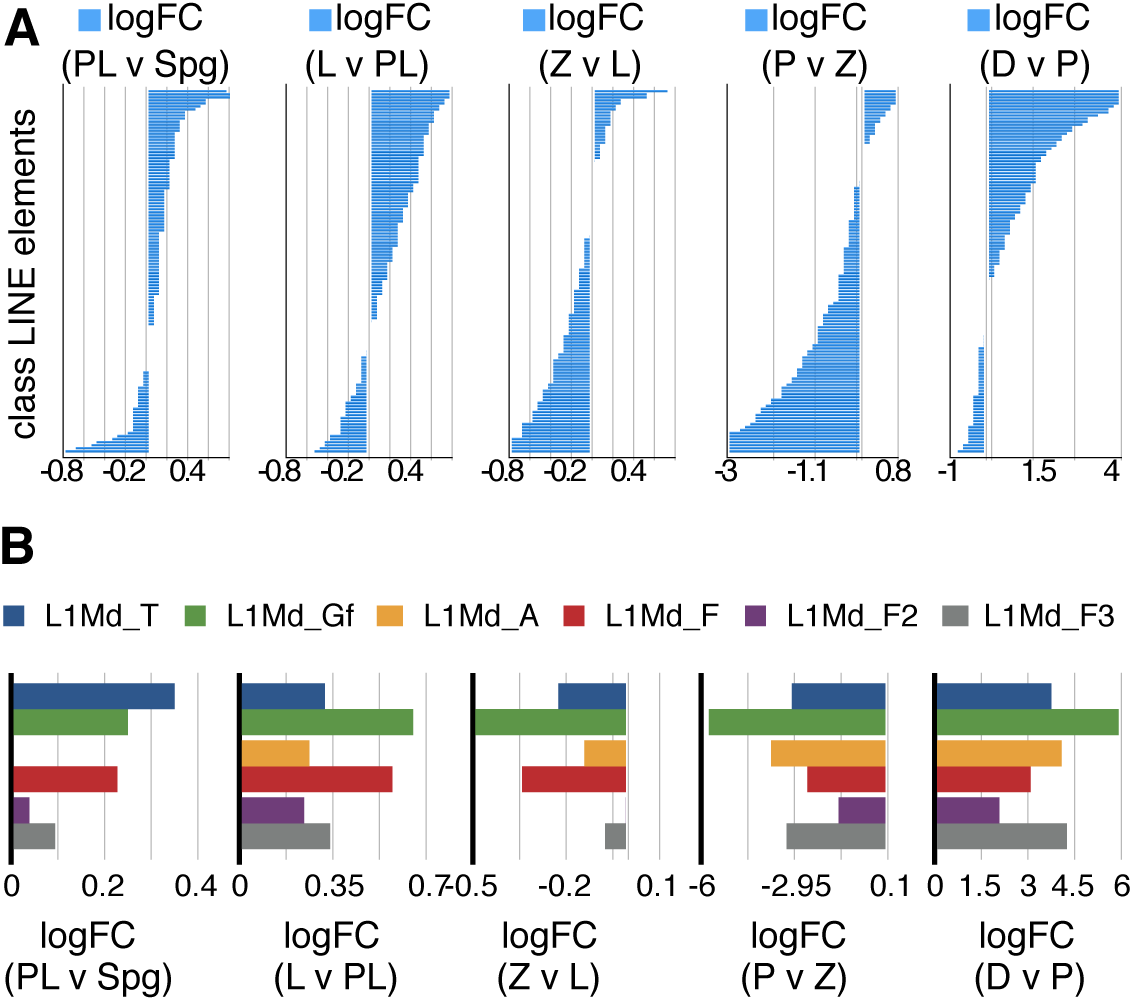
Dynamics of LINE transcript abundance in MPI. (A,B) A pairwise differential expression analysis is represented as fold change in normalized transposon counts (counts per million) between MPI stages. The horizontal bar-plot shows log2(FC) on the x-axis and a different LINE transposon from Repeat masker, on the y-axis. (A) depicts all class LINE elements, while (B) depicts select LINE-1 families that contain young and potentially active members (L1Md_T, L1Md_Gf and L1Md_A) and their progenitors (L1Md_F, L1Md_F2, and L1Md_F3).

To determine how these two bursts of L1 transcription relate to L1 protein expression, we performed immunofluorescence analysis using antibodies to the L1-encoded ORF1 protein, an acrosome-specific marker sp56, and double strand break marker γH2AX (Kim et al. 2001; Mahadevaiah et al. 2001; Soper et al. 2008). This analysis established that L1ORF1p expression in MPI begins in L, persists until mid-P and extinguishes in late P (Supplemental Fig. S13). Thus, the initial, smaller wave of L1 mRNA in the early MPI is productive while the second burst of L1 transcription at P-to-D transition does not lead to a corresponding increase in L1ORF1p levels. These L1 mRNA and protein expression dynamics fit well with the relatively low activity of the piRNA pathway in early MPI and its robust transcriptional activation in P (Di Giacomo et al. 2013; Li et al. 2013).

To conclude, in this study, we provided evidence for genome-wide TRDM at the onset of meiosis in adult male mice. Our data suggest that TRDM arises by passive, DNA replication-coupled mechanism during meiotic S phase in PL and regain of DNA methylation is a gradual process. Since cytosine methylation can occur within minutes after DNA replication, the finding of TRDM by passive DNA demethylation in pre-meiotic S phase may be viewed as unexpected (Kappler 1970; Gruenbaum et al. 1983). Yet it is consistent with diffuse nuclear localization of maintenance DNA methyltransferase DNMT1 in PL germ cells instead of its characteristic accumulation at replication foci (Jue et al. 1995). In addition, replication of 5hmC-carrying DNA could also produce hemimethylated DNA. Indeed, based on prior results of others, we do not exclude the contribution of active DNA demethylation to TRDM or later, during the P-to-D transition (Gan et al. 2013). Nevertheless, our RNA-Seq analysis of MPI cells did not support a prominent role for active DNA demethylation in PL due to the lack of *Tet1*, *Tet2*, *Aicda*/*Aid*, *Apobec1*, *Tgd* and *Smug1* expression, and very low *Tet3* transcript abundance [Supplemental Fig. S14 and (Gan et al. 2013)].

Since mouse fetal oocytes enter meiosis after genome-wide DNA demethylation, the discovery of TRDM establishes that reduced DNA methylation is a common feature both of male and female meiotic germ cells. This suggests that reduced levels of DNA methylation could be of functional significance for meiotic progression. Indeed, prior studies have shown that DNA methylation can alter meiotic recombination in fungi and plants (Maloisel and Rossignol 1998; Yelina et al. 2015b). In addition, DNA replication-coupled mechanism of TRDM suggests distinct epigenetic states of hemimethylated sister chromatids of meiotic chromosomes raising a possibility of their differential usage in MPI akin to methyl-directed mismatch repair system of *E. coli* (Putnam 2016). Finally, similarly to differential L1 expression provides the basis for the selective elimination of fetal oocytes with excessive L1 levels (Malki et al. 2014), TRDM could also contribute to gamete quality control in MPI by the unmasking deficit of DNA methylation of TEs.

## Acknowledgments

We thank Fred Tan for help with bioinformatics analyses, Svetlana Deryusheva, Safia Malki and Marla Tharp for constructive feedback on the manuscript.

## Materials and methods

### Animals

Adult C57BL/6J male mice (2-to 5-month-old) (Jackson Laboratory) were used as a source of adult testes. All experimental procedures were performed in compliance with ethical regulations and approved by the IACUC of Carnegie Institution for Science.

### Germ cell isolation

Germ cell fractions were enriched by Fluorescence Activated Cell Sorting (FACS) as described previously (Gaysinskaya and Bortvin 2015). Sorted germ cell fractions were devoid of somatic contamination but contained small amounts of germ cells from adjacent MPI stages. Cell fraction purity was determined to be >85% for Spg, ∼85% for PL, ∼85% for L, ∼80% for Z, > 90% for P, >90% for D.

### Immunofluorescence

IF on testicular sections or meiotic spreads was performed as described before (Gaysinskaya and Bortvin 2015). ImageJ was used for image analysis.

### Whole genome bisulfite sequencing (WGBS)

The protocol details are provided in the Supplemental Materials and Methods. Each biological replicate consisted of pooled cells from 2-3 different animals from different FACS procedures. For WGBS, two biological replicates (2x) were used for Spg, PL, L, Z, P, D and epididymal spermatozoa (Spz). Isolated mouse genomic DNA was spiked with approximately 0.1% unmethylated cl857 Sam7 Lambda DNA (Promega) and sheared to fragments with a range of 200-600 bp. The adaptor-ligated DNA fragments were processed for bisulfite conversion and PCR amplification. Bisulfite-treated DNA underwent 15 rounds of PCR amplification. Libraries were sequenced on Illumina HiSeq2000 platform, yielding 100bp paired-end reads. Two biological replicates were sequenced at different times. Bioinformatic analysis WGBS and DMRs is described in Supplemental Materials and Methods.

### Hairpin-Bisulfite Sequencing

Each sample consisted of pooled cells from up to two different animals from different FACS procedures. Hairpin-Bisulfite PCR was performed as previously (Arand et al), with some modifications (Supplemental Materials and Methods). Briefly, genomic DNA was digested with BspEI and the two complementary DNA strands were linked with a hairpin linker (5’P-CCGGGGGCCTATATAGTATAGGCCC). After bisulfite treatment, L1MdTf-specific PCR was performed with primers 5’TGGTAGTTTTTAGGTGGTATAGAT and 5’TCAAACACTATATTACTTTAACAATTCCCA resulting in 332-bp amplicon (Supplemental Fig. S9). The resulting product was prepared for sequencing using Ilumina *TruSeq* mRNA v2 kit, starting with end repair step, and sequenced on NextSeq 500. The 150 paired-end reads were aligned to L1MdTf promoter consensus sequence. The protocol details and bioinformatics analysis are described in Supplemental Materials and Methods.

### Accession numbers

## Author Contributions

Conceived and designed the experiments and wrote the manuscript: AB and VG. Performed the experiments: VG. Analyzed the data: VG (data analysis and bioinformatics), BM (bioinformatics and custom python scripts) and KH (bioinformatics advice and guidance related to WGBS analysis).

